# The Cilialyzer – a freely available open-source software for a standardised identification of impaired mucociliary activity facilitating the diagnostic testing for PCD

**DOI:** 10.1101/2022.09.04.506514

**Authors:** Martin Schneiter, Stefan A. Tschanz, Loretta Müller, Martin Frenz

## Abstract

Primary ciliary dyskinesia (PCD) is a rare genetic disorder causing a defective ciliary structure, which predominantly leads to an impaired mucociliary clearance and associated airway disease. As there is currently no single diagnostic gold standard test, PCD is diagnosed by a combination of several methods comprising genetic testing and the examination of the ciliary structure and function. Among the approved diagnostic methods, only high-speed video microscopy (HSVM) allows to directly observe the ciliary motion and therefore, to directly assess ciliary function. In the present work, we present our recently developed freely available open-source software – termed “Cilialyzer”, which has been specifically designed to support and facilitate the analysis of the mucociliary activity in respiratory epithelial cells captured by high-speed video microscopy.

**Methods:** In its current state, the Cilialyzer software enables clinical PCD diagnosticians to load, preprocess and replay the image sequences with a feature-rich replaying module facilitating the commonly performed qualitative visual assessment of ciliary function. The image processing methods made accessible through an intuitive user interface allow clinical specialists to comfortably compute the ciliary beating frequency (CBF), the activity map and the “frequency correlation length” – an observable getting newly introduced. Furthermore, the Cilialyzer contains a simple-to-use particle tracking interface to determine the mucociliary transport speed.

**Results:** The Cilialyzer is fully written in the Python programming language and freely available under the terms of the MIT license. The proper functioning of the computational analysis methods constituting the Cilialyzer software is demonstrated by using simulated and representative sample data from clinical practice.

**Conclusions:** The Cilialyzer serves as a useful clinical tool for standardised PCD diagnostics and provides new quantitative information awaiting to be clinically evaluated using cohorts of PCD. As the Cilialyzer is freely available under the terms of a permissive open-source license, it serves as a ground frame for further development of computational methods aiming at the quantification and automation of the analysis of mucociliary activity captured by HSVM.

## 1 Introduction

The inner surface of our airways is lined by a mucous fluid film, which is permanently propelled into the direction of the throat. The propulsion of this airway surface liquid (ASL) is provided by the collectively coordinated oscillatory motion of a myriad of subjacent cilia. Inhaled particles get entrapped by the ASL layer and subsequently transported into the direction of the throat, where they finally get swallowed. Thereby, mucociliary clearance (MCC) constitutes our airway’s primary defense mechanism by protecting our airways from inhaled toxic and infectious agents [1].

The importance of the proper functioning of motile cilia is particularly highlighted by its failure in primary ciliary dyskinesia (PCD), which is predominantly characterized by chronic airway infections. PCD is a genetic disorder with an autosomal recessive inheritance pattern and an estimated incidence of 1:10’000 to 1:20’000 [2]. Mutations in (at least) one of many genes coding for proteins involved in ciliary assembly lead to a defective ciliary structure exhibiting an impaired (or absent) ciliary motion, which in turn, is commonly thought to result in an insufficient (or absent) mucociliary clearance. The various PCD-causing genetic mutations are associated with different structural and functional defects resulting in different pathological patterns. This contributes to the challenges of diagnosis [3]. It has to be highlighted that the actual observable of interest – namely, the efficacy of the mucociliary airway clearance – is presently not directly accessible. This raises the challenging fundamental question of how to most adequately infer the efficacy of the actual mucociliary clearance and to reliably recognize an impairment of the latter.

Today, the efficacy of the actual mucociliary clearance function in PCD suspects is inferred by visually examining microscopic high-speed recordings, which merely show the ciliary motion on a few brushed nasal epithelial cells immersed in culture medium (e.g. see [4]). These conditions greatly differ from the native conditions, in which the air-exposed viscoelastic mucus layer gets propelled by the underlying collectively coordinated ciliary activity of large contiguous arrays of ciliated cells.

Due to the unavailability of computational methods allowing for a standardised analysis, the conventional HSVM analysis currently mainly relies on the visual recognition of clearly discernible defects in the ciliary beating pattern (CBP). This requires extensive empirical experience, is time-consuming and finally, is highly subjective and error-prone. The CBF is the only quantitative observable being routinely determined in diagnostic centers. Even though the conventional HSVM approach is commonly considered as the most sensitive and specific test to identify PCD, its lack of standardisation prevents its use as a confirmatory diagnostic test, or its inclusion in a diagnostic algorithm [5].

Among the currently approved diagnostic methods, however, the functional analysis performed by HSVM can be seen as the most direct method to diagnose PCD, as it allows to directly observe the mucociliary activity. Therefore, HSVM will continue to play an important role in PCD diagnostics as well as for the examination of the various functional phenotypes and the establishment of relations to the associated genotypes.

When examining the presently available body of literature, two main lines of research revolving around the development of techniques aiming towards a quantitative characterization of the high-speed recordings of ciliary activity become apparent. Motivation for both lines of research is to contribute to the standardisation of the HSVM analysis.

In the first line of research, dedicated computational methods aiming at the quantitative, objective and automated detection of defects in the ciliary motion on detached and immersed respiratory epithelial cells are developed [6–15]. Those studies mainly focus on the quantitative and automated characterization of the ciliary motion pattern of individual cilia and therefore, at making the conventional HSVM approach observer-independent. However, from a wider perspective, the reliability of the conventionally performed HSVM approach is inherently limited, as detached ciliated cells being immersed in culture medium can only partly reflect the quality of the primary mucociliary clearance mechanism. The MCC mechanism is an excellent example for a complex dynamical system with a local-to-global character, as the local fluid mechanical interactions of oscillating individual cilia are thought to generate the collective motion pattern, which eventually generates efficient mucociliary transport [16–18]. How the characteristic motion pattern of individual cilia is related to the collective ciliary motion pattern largely remains obscure. But, it can be presumed that small modulations in the individual ciliary motion pattern will amplify due to local fluid mechanical interactions and thus be more clearly reflected by the collective ciliary motion pattern [19]. This presumption is tightly connected to the second line of research, which aims to investigate and quantify the collective mucociliary phenomena in human respiratory re-differentiated cell cultures, which are grown and observed at the air-liquid interface (e.g. see [20–24]). This semi-artificial system matches the natural conditions more closely, as – in contrast to the medium-immersed brushed epithelial cells used in the classical clinical HSVM approach – presents an air-exposed viscoelastic mucus layer [25].

In summary, both lines of research have brought up new useful image processing methods allowing for a quantitative characterization of mucociliary activity. However, in diagnostics and clinical research, it is often not possible to use those image analysis routines, as either they are not freely accessible, or not provided with an intuitive easy-to-use user interface. Therefore, we have developed the herein presented freely accessible highly user-oriented open-source software, termed ‘Cilialyzer’. The Cilialyzer software is not only meant to serve as a tool for standardised diagnostic functional analyses (preferably of intact ALI cell cultures), but also as a framework for the development of further computational methods for both above-mentioned lines of research.

## 2 Methods

### 2.1 Sample preparation & imaging

The Cilialyzer software was designed for the analysis of dynamic high-speed recordings showing different types of samples from the respiratory epithelium. Therefore, neither the details of the sample preparation nor of the imaging are important for the video processing methods presented in this work. The study was approved by the ethics committees of the University Children’s Hospital Bern and of the canton Bern, Switzerland (reference no. 2018-02155). Written informed consent was obtained from every participant or her/his legal guardian. The sample preparation and the imaging used to eventually record the herein shown high-speed videos have recently been reported [26, 27]. Briefly, the image data shown here represent microscopic images showing the motion of cilia in three commonly used types of samples. Sample type I: medium-immersed human nasal epithelial cells, which were harvested by nasal brushings (see Fig. 1A). Sample type II: medium-immersed re-differentiated epithelial cells, which were scratched off from intact cultures of mucociliary epithelium grown at the air-liquid interface (ALI) (see Figure 1B). Sample type III: intact cultures of mucociliary epithelium, which were grown as well as observed at the air-liquid interface (see Fig. 1C). Note that Fig. 1A-C represent single frames of the high-speed videos (Video S1–S3), which are provided as supplemental material. After brushing or scratching, respectively, samples of type I and type II were brought into sealed chambers, which are built by a microscope slide, a 200 *μ*m thick rubber spacer and a cover glass on top. The suspensions of cells were imaged in bright field using an inverted transmitted light microscope (Olympus IX73). By using a bright field 40×-objective (Olympus UPlan Superapochromat, NA 0.95, WD 0.18 mm) an area of approximately 55×40 *μ*m^2^ was imaged onto the CMOS chip of a digital high-speed camera (GS3-U3-32S4M-C, FLIR Integrated Imaging Solutions Inc., Canada). Samples of type III were kept in culture well plates during observation. As for this kind of observation, an objective with a longer working distance was needed (Olympus UPlan SemiApo, 10× phase contrast, NA 0.3, WD 10 mm), the field of view amounts to approximately 220×165 *μ*m^2^. For all three kinds of samples, the pixel resolution was set to 640×480 and ciliary activity was recorded at 300 frames per second over a time span of 2 seconds.

**Figure 1:**
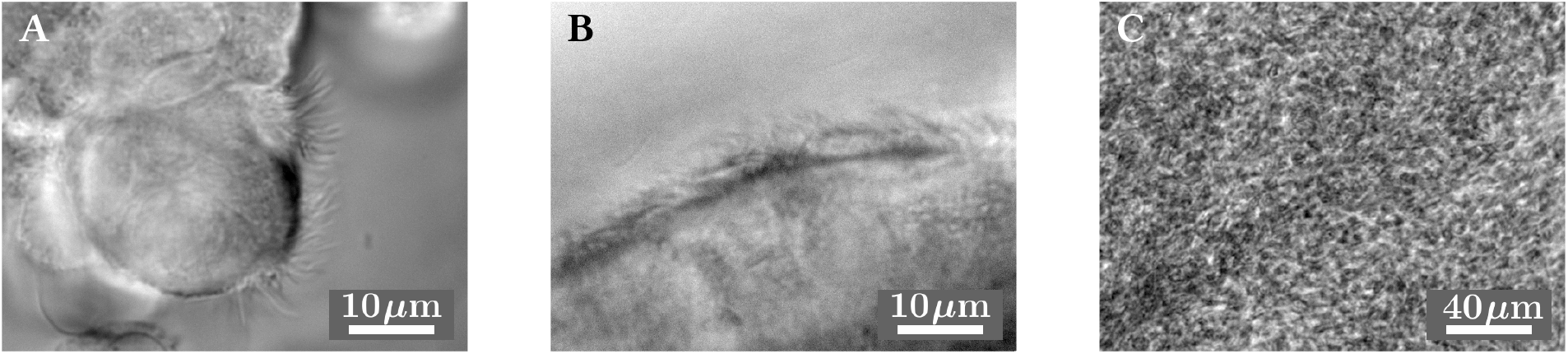
The three depicted microscopic images represent single frames of three typical high-speed recordings showing the ciliary activity within sample type I - III. The distinguishing characteristics of the different sample types can be nicely seen. A: freshly brushed human nasal epithelial cells immersed in culture medium (sample type I). This sample type is mainly characterized by the availability of only small groups of cells consisting of few ciliated cells. B: medium-immersed re-differentiated epithelial cells, which were scratched off from intact cultures of mucociliary epithelium grown at the ALI. Overall, the sample quality (in terms of cell and cilia population as well as vitality) of sample type II is typically better than in samples of type I. C: intact cultures of mucociliary epithelium grown as well as observed at the ALI (sample type III). A large array of epithelial cells can be seen (basolateral view).

For more detailed information on the sample preparation and the imaging of sample type I–III please refer to [26] and [27].

### 2.2 Video analysis procedure

Fig.2 shows an overview of the video analysis procedure for all three sample types. Each processing step is explained in detail in the following sections.

**Figure 2:**
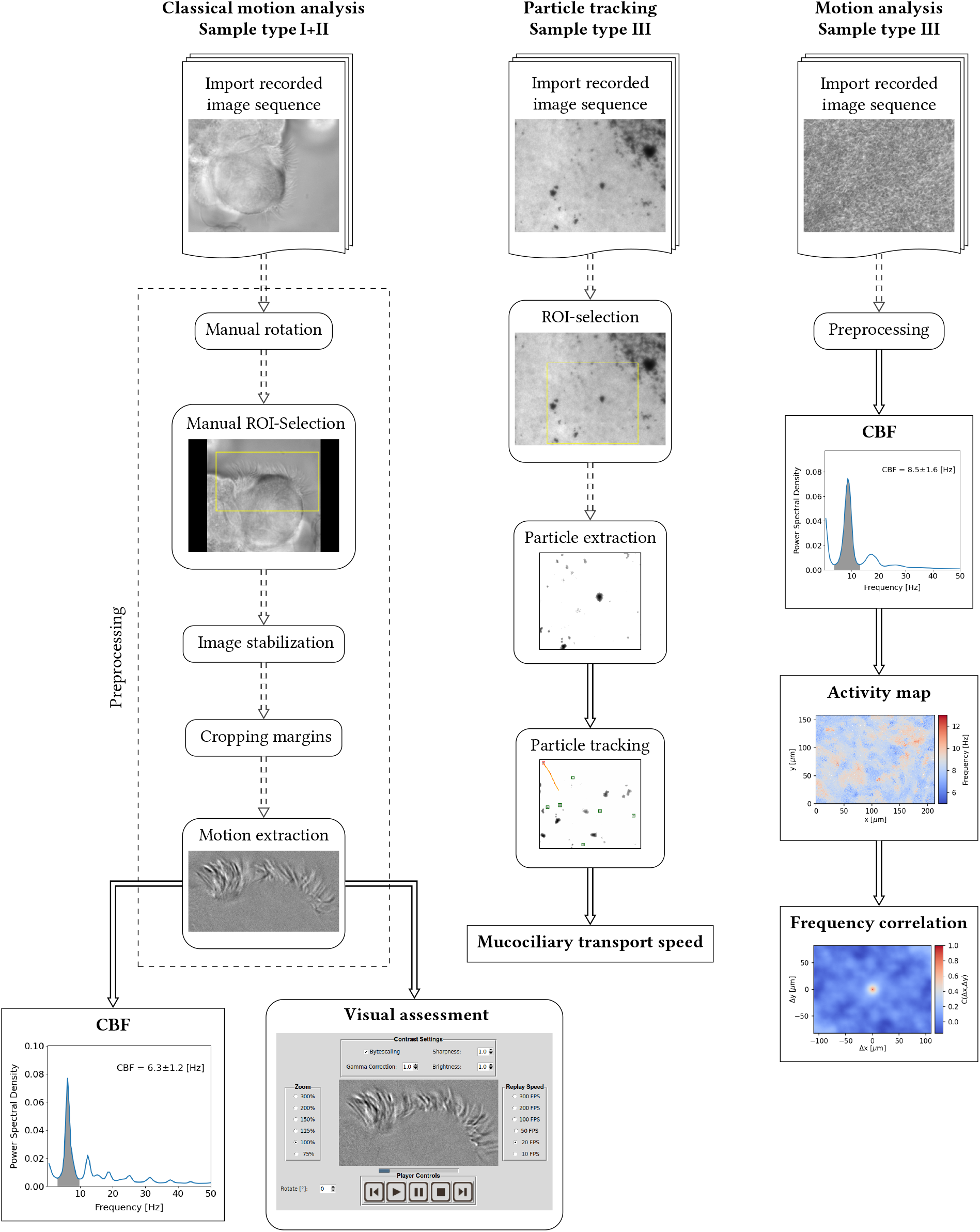
Overview of the video analysis procedures for all three sample types.

#### 2.2.1 Preprocessing

As has been shown previously (e.g. see [6, 7]), the high-speed recordings need to be adequately preprocessed before getting analyzed. In the following, we describe the available video-preprocessing steps, which mainly take care of the removal of disturbing movements and vibrations of the whole sample and the removal of the static background, emphasizing the actual signal of interest.

### Image stabilization

Image stabilization, or image registration, designates the process of aligning consecutive images and represents a highly important preprocessing step. Depending on the sample, image stabilization serves for the removal of cell or tissue motion. Before stabilizing the image sequence, the user has the possibility to manually rotate as well as to select a region of interest (ROI).

In order to get rid of undesirable sample motions, and thus to stabilize the image sequences, the Cilialyzer uses the Python package ‘pyStackReg’ [28] to align each image I [*x, y, t*] to the mean image ⟨ I [*x, y, t*] ⟩_*t*_, by performing a rigid body transform (translation and rotation). Note that ⟨…⟩_*t*_ denotes the average calculated along the time axis. Since each image is aligned to the same common reference image, the computation time could be significantly shortened by distributing the computation across *N*-1 processes running in parallel (where *N* designates the number of available CPU cores). In order to further shorten the computation time, only every second image gets actually registered and consequently, every second image is transformed in the exact same way as its predecessor.

After the image stabilization process, the image sequence needs to be slightly cropped, as the stabilization process adds invalid pixels at the margins.

### Motion extraction

As has been shown e.g. in [6, 7], the static background of an image sequence I[*x, y, t*] can be removed by subtracting the mean image:

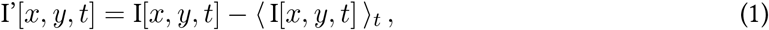

Note that the mean image subtraction, as denoted in Eq.1, is equivalent to removing the zero-frequency contribution along the time axis (for each pixel separately).

#### 2.2.2 Visual assessment

The current clinical HSVM analysis procedure is primarily based on the visual recognition of ciliary motion defects. Therefore, the Cilialyzer software was also specifically developed to improve and facilitate the visual assessment of the ciliary activity.

Clinical specialists can make use of the just described preprocessing, by: rotating the video, selecting a ROI, stabilizing the video and finally extracting the ciliary motion as needed (see Fig.3). The purpose of these preprocessing steps is to remove anything but the signal to be examined – i.e. the moving cilia. After having preprocessed the video, the Cilialyzer provides a replaying module (see Fig. 4), which allows clinicians to control the playback speed, the zoom level as well as to enhance the contrast as needed and is therefore perfectly suited for the common visual assessment of the ciliary beating pattern (e.g. see [26]).

**Figure 3:**
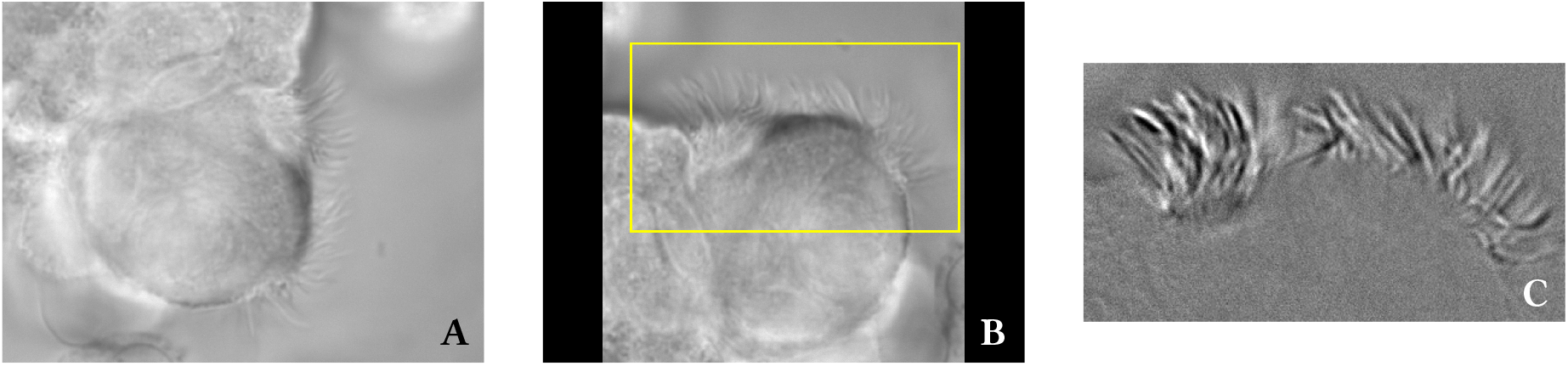
A: snapshot of an original high-speed recording showing the motion of cilia on a few epithelial cells (sample type I). The cilia can just be recognized by eye. B: The yellow surround indicates the manually selected ROI, which can be chosen in the rotated image sequence. C: snapshot of the manually selected ROI after having performed the preprocessing. The cilia as well as their motion are now more clearly distinguished from the background.

**Figure 4:**
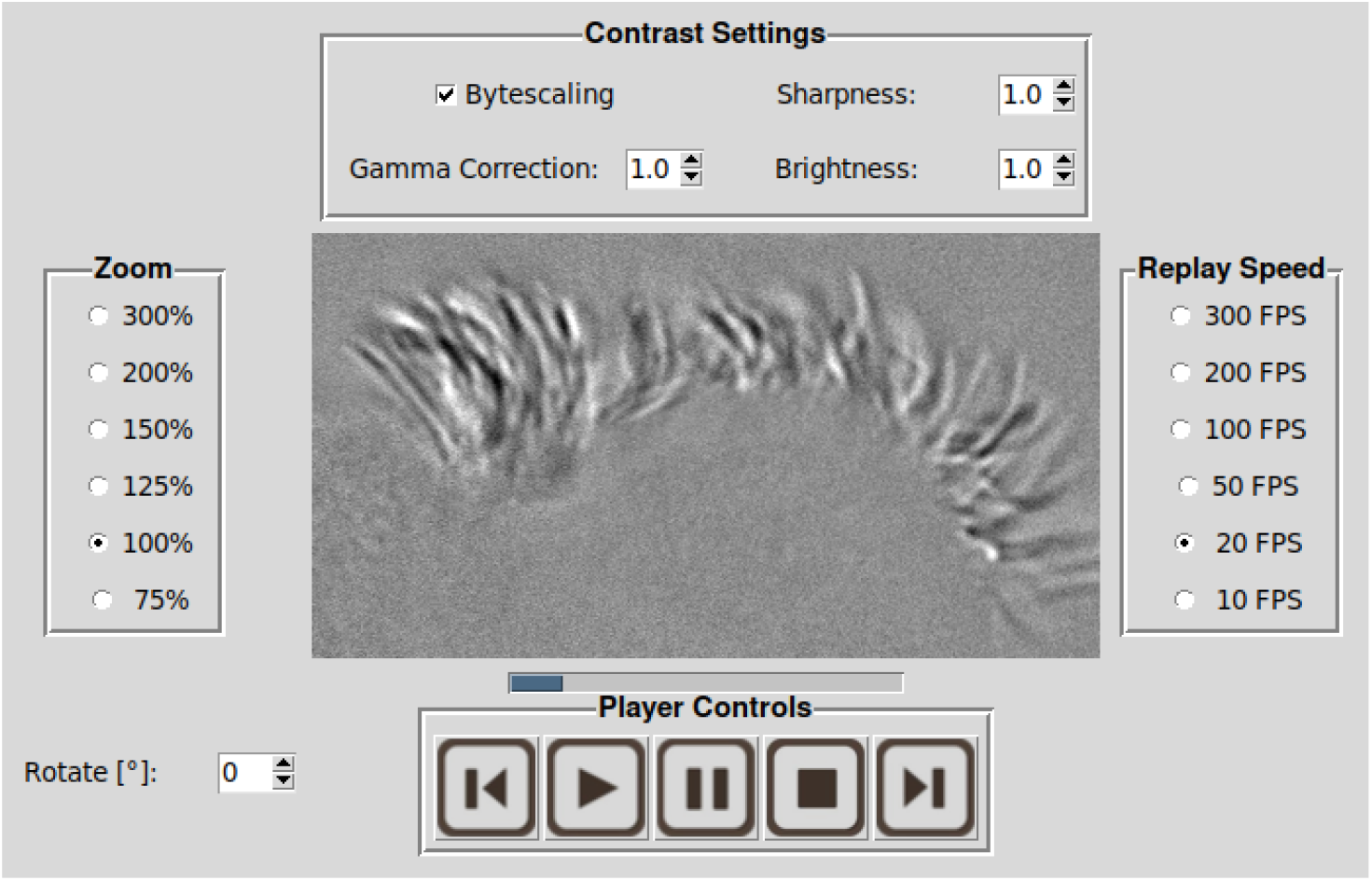
Screenshot of the Cilialyzer’s replaying module. After having extracted the ciliary motion, which is to be investigated, clinical specialists can make use of the controls, which allow to adjust the playback speed, the zoom level and the contrast.

#### 2.2.3 Computation of the ciliary beating frequency (CBF)

The ciliary beating frequency (CBF) is determined from the average power spectral density (PSD), which represents the average over all PSDs derived from the intensity variations of all the pixels within a manually selected ROI (or within the whole field of view).

In more detail, the CBF is computed as follows. The intensity variation of a single pixel located at (*x, y*) can be considered as a time series I_*xy*_(*t*). Generally, the PSD of an arbitrary time series I_*xy*_(*t*) sampled at discrete times *t* = 0, 1, …, *N*_*t*_ − 1 is defined as the absolute square of its corresponding Fourier amplitudes 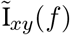, which are computed by the discrete Fourier transform:

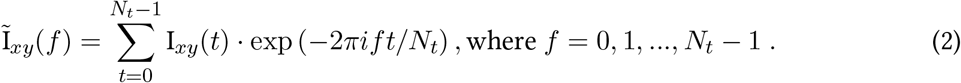

The discrete Fourier transform (Eq. 2) was computed by using the highly efficient Fast Fourier transform (FFT) [29]. The PSD of a single pixel located at (*x, y*), denoted by P_*xy*_(*f*), is finally given by the absolute square of its complex Fourier amplitudes: 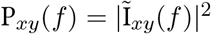. In order to get insight into the frequency contributions within the manually selected ROI (or the whole field of view), comprising *N*_*x*_ · *N*_*y*_ pixels, the Cilialyzer eventually displays the average PSD, denoted as ⟨ P(*f*) ⟩_*xy*_, in the positive frequency range:

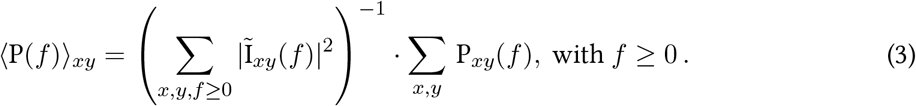

Please note that the displayed average power spectral density gets slightly smoothed with a gaussian kernel G_*σ*=1_, which can be denoted by the following convolution: ⟨ P(*f*) ⟩_*xy*_ ∗ G_*σ*=1_. Furthermore, the zero-frequency contribution gets omitted.

The CBF is finally determined as the PSD-weighted average within the band-width belonging to the main peak, which is indicated by the gray shading in Fig.5. Note that the series of peaks at integer multiples of the CBF represent higher harmonics, which indicate that the intensity variations are not purely harmonic. The calculation of the CBF requires two cut off frequencies *f*_1_ and *f*_2_, which are given by the lower and upper limit of the main peak in the average PSD. The Cilialyzer automatically suggests those two cut off frequencies (according to the procedure explained in Sec. S1). However, the user has also the possibility to manually adjust the bandwidth selection.

**Figure 5:**
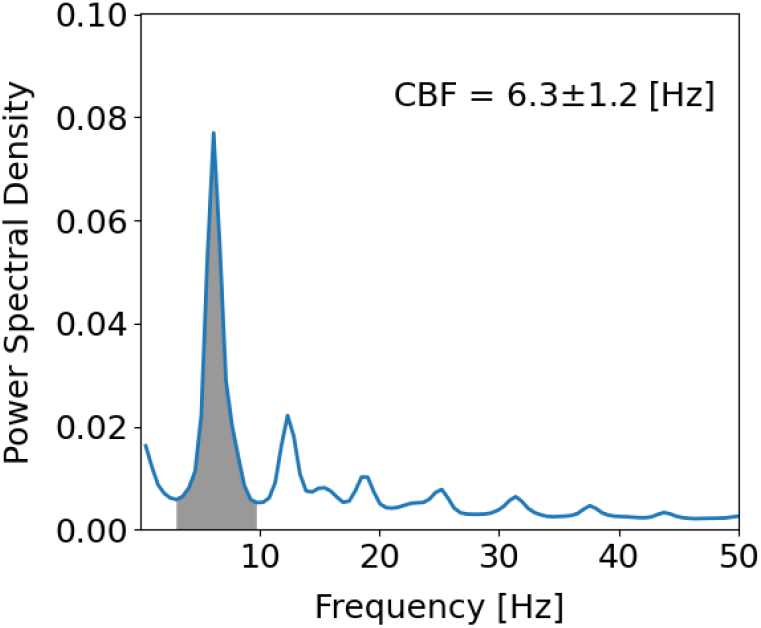
Typical power spectral density (PSD) computed for the rectangular user-selection shown in Fig.3B.

The mean CBF, denoted as ⟨*F*⟩ and the peak width, which is measured by the weighted standard deviation *σ*_*F*_, are then calculated according to:

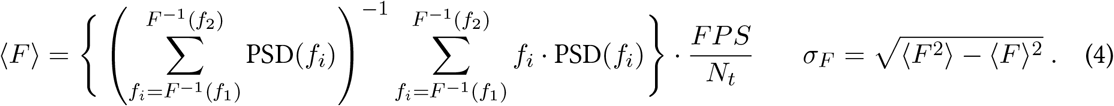

#### 2.2.4 Activity map

The activity map provides information on how the CBF is spatially distributed and can thus be seen as a position-resolved CBF map. The computation of the activity map follows the principles, which have previously been presented [30].

Let us denote the pixel-specific power spectral density as PSD_*xy*_(*f*). Usually, only a part of the field of view displays ciliary activity. Therefore, in order to discriminate between pixels exhibiting a reasonable PSD from invalid pixels, we make use of the following simple condition. On the basis of the average PSD, we can define the total power *P* contributed by all the frequencies found inside the bandwidth of the main peak 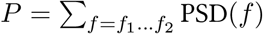, where PSD(*f*) denotes the area averaged power spectral density. Furthermore, let *P*_*xy*_ be the pixel-specific total power contributed in the bandwidth of the main peak, i.e. 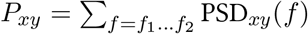. Finally, a pixel located at (*x, y*) is marked as invalid (NaN), if: *P*_*xy*_*/P* < Θ, where Θ ∈ (0, 1) represents a threshold, which can be adjusted manually. For valid pixels, on the other hand, a location-resolved CBF is ascribed, which is computed according to Eq.4 by replacing the area averaged PSD with the pixel-specific PSD, i.e. PSD_*xy*_(*f*).

#### 2.2.5 Frequency correlation

The ‘frequency correlation’, denoted by *ξ*, refers to the correlation length in the spatial autocorrelation of the activity map and constitutes another feature of the Cilialyzer. The spatial autocorrelation of the activity map gets computed by exploiting the Wiener-Khinchin theorem, which states that the correlation function and the power spectrum are a pair of Fourier transforms. The inverse Fourier transform of the spatial power spectrum therefore yields the autocorrelation function. As the computation in the Cilialyzer takes the zero-padding as well as missing values into account (i.e. pixels showing an invalid power spectral density), the computation of the spatial autocorrelation of the activity map is not trivial. The interested reader is therefore referred to Sec.S2, where the computational details are outlined.

In simple terms, the frequency correlation *ξ* can be understood as the average distance over which the CBF roughly remains preserved. Another interpretation concerns *ξ*^2^, which can be seen as the average patch size over which the frequency roughly remains preserved. Those patches can clearly be recognized by eye in the activity map provided in Fig.6.

**Figure 6:**
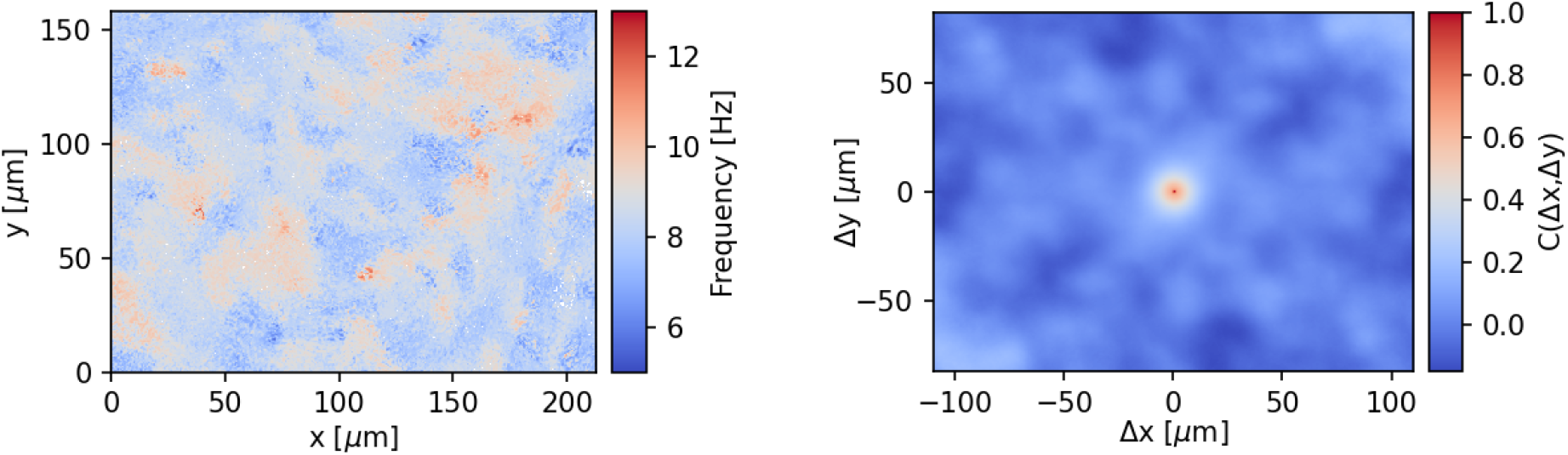
Left: activity map showing the spatial distribution of the CBF in a re-differentiated cell culture grown at the air-liquid interface. Right: frequency correlation i.e. the spatial autocorrelation of the activity map shown on the left. The size of the central peak can be seen as a measure for the average patch size recognizable in the activity map.

#### 2.2.6 Mucociliary transport speed (particle tracking)

Especially in samples of type III, mucociliary transport represents a potentially insightful observable providing rather direct information on the efficacy of the mucociliary clearance mechanism. In order to adequately track the movement of the mucus layer, it is usually necessary to apply some kind of tracers. Please note that particle suspensions – even though frequently used – are not suited for the investigation of mucociliary transport, as droplets applied on the outermost mucus layer change the physical properties of the air-liquid interface and might even induce transport phenomena, which are not related to the underlying ciliary propulsion mechanism. Therefore, we recommend to blow a tiny amount of lightweight dry puff ball spores (from calvatia excipuliformis) onto the mucus surface, as was done previously [27, 30–33].

The Cilialyzer’s particle tracking module consists of a simple-to-use interface, which allows the user to extract and select the particles to be tracked. Since the tracers are small point-like structures, they can optionally be extracted by subtracting a basically blurred copy (Lee-filtering followed by Gaussian filtering) from the original image sequence. A subsequent thresholding removes some residual background. The particle tracking is based on a local extremum tracker within a manually adjustable window size. This yields noisy particle trajectories, which need to be adequately fitted, in order to correctly determine the curve length. As has been observed previously in vitro [21, 24], mucociliary epithelium grown at the ALI in circularly confined culture vessels typically exhibit circular mucociliary transport. This means that particles can run along circular, spiral or quasi-linear trajectories. Therefore, the noisy particle traces are first slightly smoothed and subsequently fitted by a univariate spline, whose curve length is finally used to determine the mucociliary transport speed. This procedure is highly robust and can cope with arbitrary particle trajectories.

## 3 Results

### 3.1 Software

The herein presented computational methods constitute the Cilialyzer, which is a freely available open-source software. It is fully written in the Python programming language [34] and distributed under the terms of the MIT license with the additional request to include a citation of the present manuscript. The source code, tutorials as well as compiled releases are provided at Github: https://github.com/msdev87/Cilialyzer. Readers who are only interested in downloading and using the application are referred to: https://msdev87.github.io/Cilialyzer.

### 3.2 Preprocessing

On the one hand, an adequate preprocessing improves the visual assessment, as the image stabilization and the subsequent subtraction of the static background highlights the moving cilia to the observer. On the other hand, distorting movements of the whole sample must be eliminated in order to ensure the correctness of the objective quantitative analysis (see the example of a compromised CBF computation in the subsequent section). The enclosed videos S4–S6 are meant to illustrate the power of the presently available preprocessing and represent preprocessed versions of videos S1–S3.

### 3.3 Validation of the CBF-computation

It has been shown repeatedly (e.g. see [35, 36]) that the CBF computed by means of the power spectral density of the pixel’s intensity variations corresponds well to the manually determined CBF, which is performed by progressing the high-speed videos frame by frame in order to visually count the ciliary oscillation period. Therefore, we decided to not show this agreement again. However, similar to the approach chosen in [37], the correctness of the output frequency can be shown by recovering a predefined input frequency. For this purpose, we have simulated a metachronally coordinated array of cilia, which is shown in Fig. 7. See also video S7, which provides an example of the ciliary array oscillating at 10 Hertz. This array of cilia was used to generate 20 videos, in which the cilia oscillate with a predefined input oscillation frequency ranging from 1 to 20 Hertz. Fig. 8 shows that the input frequency (simulated CBF) corresponds exactly to the output frequency (measured CBF) determined by the Cilialyzer.

**Figure 7:**
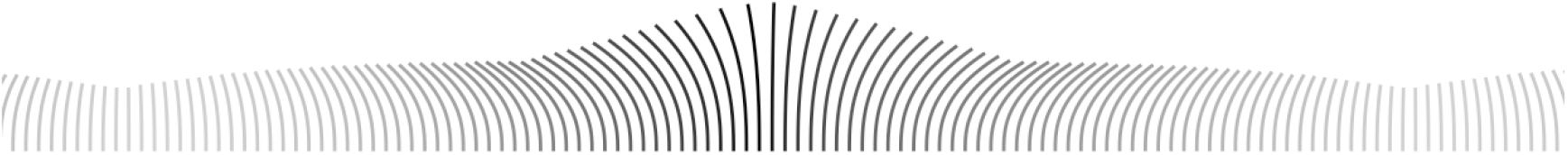
Snapshot of the simulated ciliary array used to validate the computation of the CBF.

**Figure 8:**
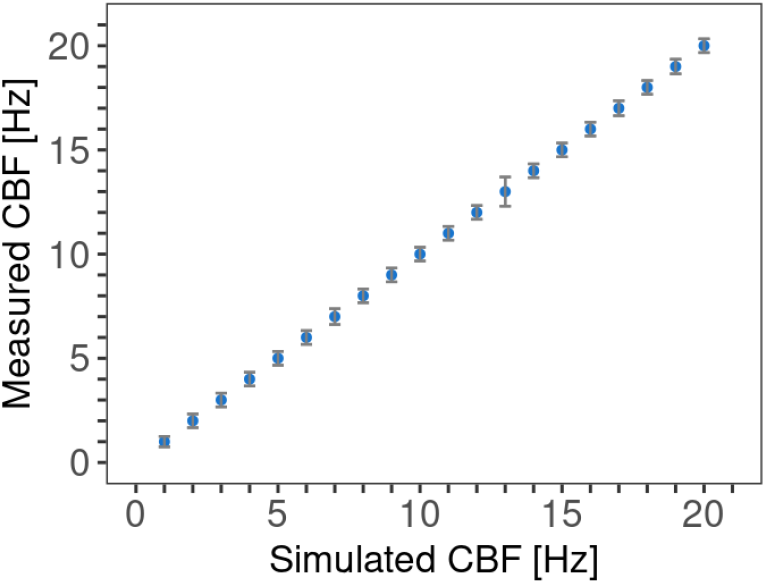
For the functional testing of the CBF-computation, an array of cilia was simulated to oscillate in the range from 1 to 20 Hertz, which corresponds to the input frequency shown on the x-axis. The graph shows that the CBF as determined by the Cilialyler (shown on the y-axis) exactly matches the simulated input frequency.

#### 3.3.1 Importance of the image stabilization

The important role of the image stabilization is best illustrated with an example from clinical practice. When considering the high-speed recording provided by video S8, which shows a freshly excised group of nasal epithelial cells (sample type I), it can be seen that the ciliary motion is superposed by a quasi-periodic movement of the whole group of cells. Video S9 shows a stabilized and cropped version of video S8. Fig. 9A shows the CBF computation based on the non-preprocessed video S8, whereas Fig. 9B shows the CBF computation based on the preprocessed video S9. In Fig. 9A, it can be nicely seen that without any preprocessing the average power spectrum gets completely compromised by the superposed motion of the whole group of cells. The actual CBF-peak is hardly recognizable and the selection of the CBF-bandwidth becomes a guessing game. Whereas, the average power spectral density we are actually looking for and which corresponds to the ciliary motion can be recovered by preforming the preprocessing, as can be seen in Fig. 9B.

**Figure 9:**
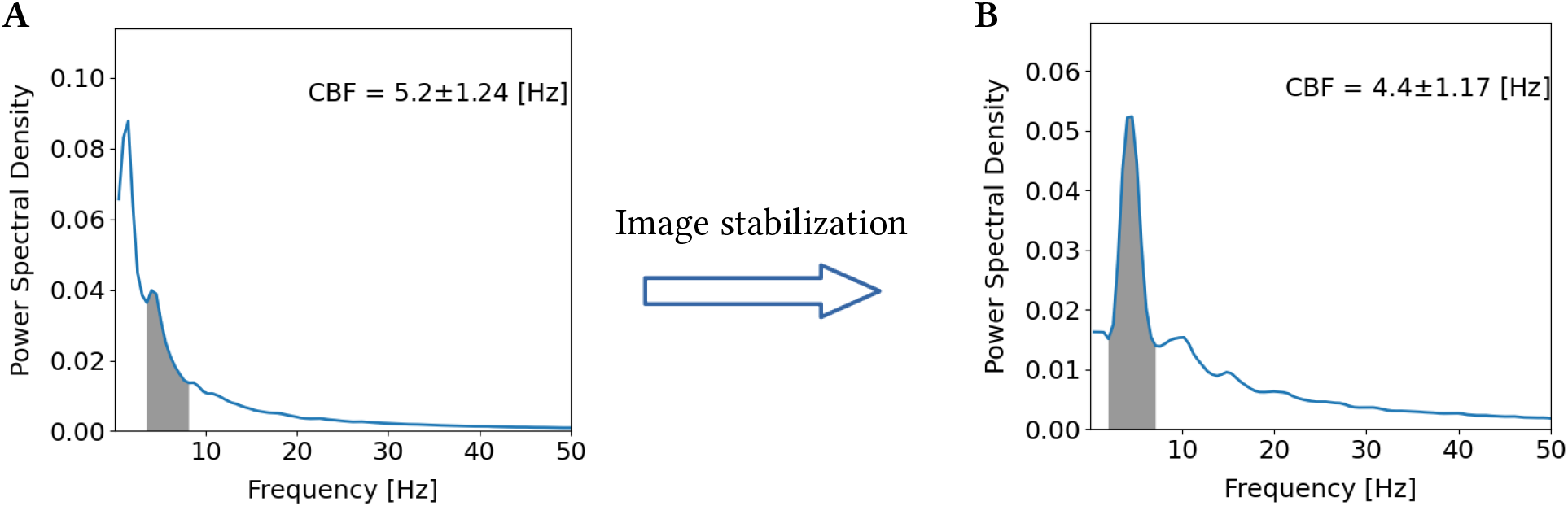
A: average power spectral density based on the non-preprocessed video S8: the actual CBF-peak is hardly recognizable and the selection of the CBF-bandwidth becomes a guessing game. B: After performing the preprocessing, the average power spectral density, which actually corresponds to the ciliary motion becomes apparent.

### 3.4 Spatial distribution of the CBF

Similar to the validation of the CBF-computation presented in the previous section, we verified the proper functioning of the computation of the activity map and the frequency correlation length with simulated data.

In order to illustrate the proper functioning of the methods, the analysis of a single example video used for functional testing is shown in the following. Fig. 10A represents a snapshot of the simulated image sequence, which is provided in the supplementary material (see video S10). The artificially generated video shows the propagation of harmonic plane waves, for which the temporal frequency was chosen differently and randomly in the range between 10 and 20 Hertz for 10×10 regions of interest - which is correctly reproduced and visualized by the activity map shown in Fig. 10B. Fig. 10C finally shows the spatial autocorrelation function of the activity map and displays the frequency correlation length *ξ*_*f*_. In this particular example, a pixel resolution of 400×400 pixels was chosen. Furthermore, the frequency was chosen to be constant within each ROI consisting of 40×40 pixels. In the Cilialyzer we assumed that one pixel measures 1 *μ*m^2^. Considering that the frequency correlation is intended to represent an observable, which provides rather relative ^1^ than absolute information about the distance over which the CBF roughly remains preserved, the absolute value of roughly 44 pixels is very close to the actual ROI size, which was chosen to be 40 pixels.

**Figure 10:**
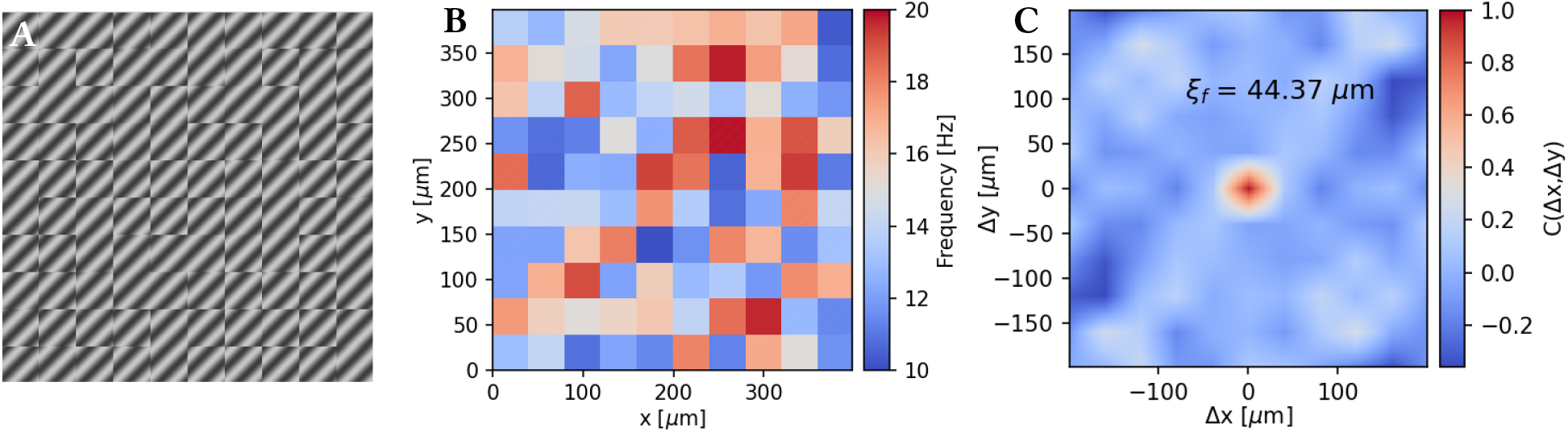
For the functional testing of the computation of the activity map and the frequency correlation length, we simulated in particular harmonic plane waves (displayed in the left panel) with temporal frequencies ranging from 1 to 20 Hertz, as is shown by the activity map shown in B. C: The frequency correlation length *ξ*_*f*_ provides a measure for the distance over which the frequency remains roughly preserved.

## 4 Discussion

This work presents the current version of the freely available Cilialyzer software, which we specifically developed to improve and facilitate a standardised analysis of the ciliary function captured by HSVM. We envision the Cilialyzer software not only as a clinical tool for the analysis of mucociliary activity, but also as a ground frame for the future development of an even more comprehensive consensus-based application, which allows clinicians as well as biomedical researchers to automatically and quantitatively detect impaired mucociliary activity. The functional testing, i.e. the proper implementation of the accessible computational methods, is demonstrated by using either simulated data or examples from clinical practice. Therefore, the Cilialyzer software can be considered validated and used clinically with immediate effect.

In comparison to applications, which were developed for the analysis of ciliary function in HSVM recordings and were recently made publicly available (e.g. see [35, 36, 38]), the Cilialyzer software offers many advantages. The present version can be easily customized and extended, as Python is well suited for rapid prototyping. Users do not need a license or a virtual machine to run the software, and for users working on Microsoft Windows, there is the possibility to download a compiled binary release, which is available on GitHub: https://github.com/msdev87/Cilialyzer. The Cilialyzer represents the most comprehensive application serving for the analysis of mucociliary activity in HSVM recordings. From a clinical point of view, the Cilialyzer stands out not only because of its ease of use, but also because of the many pre-processing and replaying options it offers. After loading an image sequence (in any common image format), the Cilialyzer particularly offers the possibility to easily rotate the sequence, select a ROI, remove undesirable tissue motion, remove the static background, enhance the contrast, zoom-in and set the playback speed. The enclosed videos S4-S6 and S9, which represent pre-processed versions of the videos S1-S3 and S8, respectively, are intended to provide an idea of the available pre-processing options. In the pre-processed videos, the ciliary motions are clearly emphasized, which supports and facilitates the common visual assessment of the ciliary beating pattern. In recordings captured by HSVM, the ciliary motion is often superposed by motions of the whole specimen or by motions of the cells (see video S8). As shown in Sec. 3.3.1, such videos can be stabilized (see video S9) allowing for a reliable CBF-determination.

For the above-mentioned reasons, the Cilialyzer is ideally suited for the current clinical assessment of ciliary function in medium-immersed epithelial airway cells (sample type I and II), which primarily consists of the CBF-computation and the visual assessment of the ciliary motion pattern (see [26]).

Thanks to the availability of affordable high-performance computers and the ongoing development of new dedicated image processing techniques, it will soon be possible to retrieve considerably more quantitative information from the high-speed recordings. We are strongly convinced that this will significantly enhance the reliability of future HSVM outcomes in PCD diagnostics.

From a wider perspective, detached ciliated cells being immersed in culture medium can only partly reflect the quality of the primary mucociliary clearance mechanism, which is the actual variable of interest to be judged. We therefore believe that specimens of sample type III, i.e. intact re-differentiated respiratory cell cultures, which are grown as well as observed at the air-liquid interface, harbor the greatest potential in terms of the information content about the primary mucociliary clearance. The following reasons support this presumption. 1) In contrast to brushed respiratory epithelial cells suspended in culture medium, cell cultures grown at the air-liquid interface match the natural conditions more closely, as they present an air-exposed mucus layer. 2) The actual variable of interest, namely the mucociliary transport speed, can be measured directly after applying lightweight dry puff ball spores, which have been used previously [27, 30–33] and are ideally suited to track the movement of the mucus layer. 3) Compared to the recordings of medium-immersed brushed epithelial cells, the recordings of ciliary activity in intact ALI cultures are much more homogeneous – in terms of their occurrence and orientation to the optical axis – facilitating the use of quantitative image processing methods. 4) The ciliary activity of a much greater number of cells can be recorded. Since we know that the CBF can vary significantly between individual cells – or between groups of cells, averaging over a much greater number of cells is expected to significantly increase the diagnostic value of the CBF. 5) Finally, mucociliary transport represents a collective phenomenon and it can be presumed that small modulations in the individual ciliary motion pattern will amplify due to local fluid mechanical interactions and thus be more clearly reflected by the collective ciliary motion pattern [19].

Therefore, we are confident that the HSVM analysis will run in the direction of this newly emerging type of genuine functional PCD diagnostics and have started to develop mathematical methods allowing for the quantitative (and automated) characterization of the collective mucociliary activity in ALI cultures. Currently, the Cilialyzer allows to determine the CBF, the mucociliary transport speed and the frequency correlation length for videos recorded in any open format on any microscope equipped with a digital high speed camera. The value of these three observables for PCD diagnostics, as determined in recordings showing the mucociliary activity in cell cultures, remains to be properly studied. The CBF, the mucociliary transport speed as well as the frequency correlation length can be considered as validated and can be used for clinical studies with immediate effect.

Future incorporations of further methods, will allow for a more accurate characterization of the collective ciliary motion in ALI cultures, which in turn, will allow for a more sensitive identification of impairments of the collective mucociliary activity. For clinicians, we therefore strongly recommend to use not only a PCD analysis software – like our Cilialyzer, which provides quantitative information about ciliary function, but as important, ALI cultures grown according to a standardised protocol.

Finally, we are looking forward to a broad user and developer community and hope that with the help of the Cilialyzer software, many of the mentioned open questions will be addressed in clinical studies.

## Acknowledgments

This work was co-funded by the Swiss Lung Foundation. Part of the microscopic work was performed on equipment of the Microscopy Imaging center (MIC) of the University of Bern. Study authors participate in the BEAT-PCD clinical research collaboration (CRC), supported by the European Respiratory Society. Our PCD-UNIBE diagnostic center participates in the European Reference Network (ERN) Lung PCD core as a supporting member.

## Ethical approval

The study was approved by the ethics committees of the University Children’s Hospital Bern and canton Bern, Switzerland (reference no. 2018-02155). Written informed consent was obtained from every participant or her/his legal guardian.

## Author Contributions

**Martin Schneiter:** Conceptualization; Data curation; Formal analysis; Investigation; Methodology; Software; Validation; Visualization; Writing – original draft preparation; Writing – review & editing. **Stefan A. Tschanz:** Conceptualization; Funding acquisition; Investigation; Methodology; Project Administration; Resources; Supervision; Validation; Writing – original draft preparation; Writing – review & editing. **Loretta Müller:** Conceptualization; Funding acquisition; Investigation; Methodology; Project administration; Resources; Writing – review & editing. **Martin Frenz:** Conceptualization; Funding acquisition; Investigation; Methodology; Project administration; Supervision; Writing – review & editing.

## Supplementary Materials for

### S1 Automated CBF-bandwidth selection

As is explained in the main text, the Cilialyzer displays the ROI-related, gaussian-smoothed average power spectral density ⟨P(*f*)⟩_*xy*_ ∗ G_*σ*=1_ from which the zero-frequency contribution is omitted. The bandwidth of cilia-generated frequency contributions, i.e. the peak width of the fundamental frequency, is then found as follows. First, the function ⟨P(*f*)⟩_*xy*_ ∗ G_*σ*=1_ (for *f >* 0) is fitted by the fit function *g*(*f*), which is composed of a decaying power function and two Gaussians:

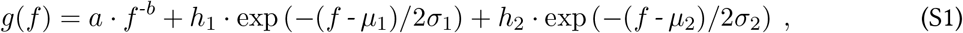

where all the fit parameters *a, b, h*_1_, *μ*_1_, *σ*_1_, *h*_2_, *μ*_2_ and *σ*_2_ are ≥ 0. The power function fits the ‘background’, which represents so-called 1/f noise being present in many biophysical systems. In some cases, a single Gaussian is not sufficient to fit the fundamental frequency, which is why the sum of two Gaussians is fitted. Before proceeding, the algorithm first checks, whether the second Gaussian is indeed different from the first one. In the case that *μ*_2_ lies outside of the interval [*μ*_1_ − *σ*_1_, *μ*_1_ + *σ*_1_], the two Gaussians are treated as being different. As the average PSD typically exhibits higher harmonics, it must then be checked, whether the second Gaussian contributes to the fundamental frequency, or, represents a higher harmonic. If *μ*_2_ + *σ*_2_ < 2 · *μ*_1_, the second Gaussian is not treated as a higher harmonic and is treated as to represent a contribution to the fundamental frequency. The CBF-bandwidth is finally selected depending on whether the second Gaussian represents a higher harmonic or a contribution to the fundamental frequency. In the latter case, *f*_1_ is given by the minimum value of the PSD in the interval [*μ*_1_ −4*σ*_1_, *μ*_1_ −2*σ*_2_] and *f*_2_ as the minimum value of the PSD in the interval [*μ*_2_ +*σ*_2_, *μ*_2_ +4*σ*_2_], if *μ*_1_ < *μ*_2_ or, if *μ*_1_ *> μ*_2_ then *f*_1_ is given by the minimum PSD value in [*μ*_2_ − 4*σ*_2_, *μ*_2_ − 2*σ*_2_] and *f*_2_ by the minimum PSD value in [*μ*_1_ + *σ*_1_, *μ*_1_ + 4*σ*_1_] .

### S2 Computation of the frequency correlation

This section specifies the algorithm to determine the frequency correlation length *ξ* implemented in the Cilialyzer.

Let us denote the activity map by **A**[*x, y*], which holds the position-specific CBF for each pixel located at (*x, y*) and where *x* ∈ {1, …, *N*} and *y* ∈ {1, …, *M*}. Furthermore, **M**_**v**_[*x, y*] represents a binary validity-mask, which indicates whether a valid frequency could be assigned to each pixel *x, y*. In other words, missing values in the activity map are marked as: **M**_**v**_[*x, y*] = 0. In a first step, the activity map has to be centered by subtracting its mean value:

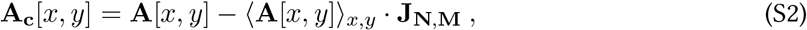

where **A**_**c**_[*x, y*] denotes the centered activity map and **J**_*N,M*_ the *N* × *M* matrix containing just ones. The mean value of the activity map is calculated by ignoring missing values, i.e. pixels showing an invalid power spectral density. This can be denoted as follows:

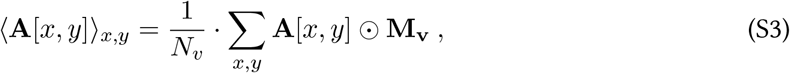

where the operator ‘0’ denotes the element-wise product and *N*_*υ*_ = ∑ _*x,y*_ **M**_**v**_ the number of valid pixels.

In a next step, we pad the centered activity map **A**_**c**_ in such a way that the zero-padded activity map, which we denote by **A**_**c**+**0**_, is twice the size along each dimension, i.e. 2*N* × 2*M*. The zero-padding can be denoted as follows:

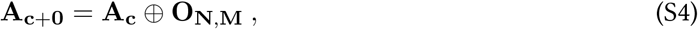

where ‘⊕’ denotes the direct sum and **O**_**N**,**M**_ the zero matrix with dimension *N* × *M*. By using the Wiener-Khinchin theorem, we compute the ‘pseudo-autocovariance’, which is biased due to the zero-padding as well as possible missing values:

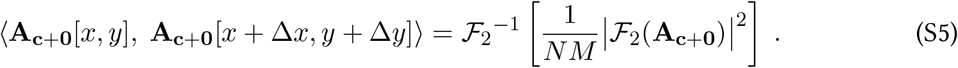

In Eq.S5, ⟨**A**_**c**+**0**_[*x, y*], **A**_**c**+**0**_[*x* + Δ*x, y* + Δ*y*]⟩ denotes the pseudo-autocovariance. ℱ_2_ and 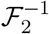 the two-dimensional Fourier transform and its inverse, respectively.

In a further step, we need to correct for the zero-padding as well as for the missing values (which are set to zero), which can be done by repeating the processes specified by Eq.S4 and Eq.S5 for the validity-mask **M**_**v**_. This delivers the normalization factors, which are given by:

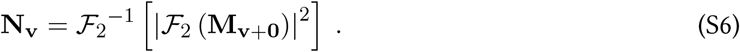

The ‘error’ introduced by the zero-padding as well as by possible missing values can now be corrected by an element-wise division of the pseudo-autocovariance by **N**_**v**_. This yields the autocovariance **C**′[Δ*x*, Δ*y*]:

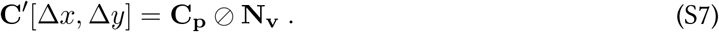

An element-wise division of the autocovariance by the variance in **A**_*c*+0_, yields the two-dimensional autocorrelation of the activity map **C**[Δ*x*, Δ*y*].

The frequency correlation length *ξ* is finally determined by counting the number of pixels in the correlogram **C**[Δ*x*, Δ*y*], for which the correlation is greater than *e*^*−*1^. Those pixels – multiplied by the actual pixel size in real units – represent the average patch size *ξ*^2^ over which the frequency roughly remains preserved.

### S3 Supplementary videos

All videos showing ciliary activity on real cellular material were recorded at 300 FPS over a time span of two seconds.

**Video S1:** The video represents a non-preprocessed recording captured by HSVM and shows the ciliary activity of a few medium-immersed ciliated cells, which were derived by a nasal brushing in a healthy volunteer. Freshly brushed cell material is characterized by a poor amount of ciliated cells, which are usually only available in the form of small groups of cells consisting of very few ciliated cells. The video is encoded with a framerate of 50 FPS (6× slower than real time).

**Video S2:** The video represents a non-preprocessed high-speed recording of the ciliary activity on medium-immersed re-differentiated epithelial cells, which were scratched off from intact cultures of human nasal mucociliary epithelium, which were grown at the air-liquid interface. The video is encoded with a framerate of 50 FPS (6× slower than real time).

**Video S3:** The video represents a non-preprocessed HSVM recording of an intact culture of mucociliary epithelium, which was grown at the ALI. During recording, the culture was kept within a closed culture well plate. As for this sample type, the imaging is done through the light-scattering mucus layer, the collective (muco)ciliary activity within the culture is hardly recognizable without any preprocessing. The video is encoded with a framerate of 50 FPS (6× slower than real time).

**Video S4:** This video represents a preprocessed version of video S1. The preprocessing consists of rotating the images, a ROI-selection, the image stabilization, the subtraction of the mean image, the byte-scaling, a slight contrast enhancement (*γ* = 1.2) and a magnification of the images (zoom level of 150%). The video is encoded with a framerate of 30 FPS (10× slower than real time).

**Video S5:** represents a preprocessed version of video S2. The following preprocessing was applied: video S2 was rotated, a selected ROI was stabilized, slightly zoomed (125%) and byte-scaled (no mean subtraction and no contrast enhancement – besides byte-scaling – was applied). The video is encoded with a framerate of 30 FPS (10× slower than real time).

**Video S6:** This video represents a preprocessed version of video S3. In this case, the preprocessing only consists of the mean image subtraction and a slight contrast enhancement. In contrast to its non-preprocessed version, one can clearly observe the collectively coordinated ciliary activity. The video is encoded with a framerate of 30 FPS (10× slower than real time).

**Video S7:** The video shows a simulation of an antiplectically coordinated array of cilia. The video is encoded with 10 FPS and shows the motions of the cilia as if they would oscillate at 10 Hertz and were recorded at 300 FPS.

**Video S8:** The video shows a freshly brushed group of nasal epithelial cells (sample type I). Besides byte-scaling, no preprocessing was applied. It can be seen that the ciliary motions are superposed by a quasi-periodic motion of the whole group of cells. The video is encoded with a framerate of 50 FPS (6× slower than real time).

**Video S9:** This video represents a preprocessed version of video S8. The preprocessing consists of rotating the images, a ROI-selection, the image stabilization, the subtraction of the mean image, the byte-scaling, a slight contrast enhancement (*γ* = 1.2) and a zoom. The video is encoded with a framerate of 30 FPS (10× slower than real time).

**Video S10:** The video shows a simulation of propagating harmonic plane waves with temporal frequencies ranging from 10 to 20 Hertz. The video is encoded with 20 FPS and shows the motion of the plane waves as if they were captured at 100 FPS.

## Footnotes

1 As compared to recordings captured according to the same experimental procedure.

